# How does it feel to be on psilocybin? Dose-response relationships of subjective experiences in humans

**DOI:** 10.1101/2020.06.09.142802

**Authors:** Tim Hirschfeld, Timo Torsten Schmidt

## Abstract

Psilocybin is the active component of magic mushrooms and is well known for its psychoactive properties. Different questionnaires have been developed to systematically quantify altered states of consciousness induced by psychoactive drugs. The aim of this study was to obtain the dose-response relationships of the subjective experiences induced by psilocybin in healthy study participants. For this purpose, we applied a linear meta-regression approach on questionnaire ratings after oral administration of psilocybin in a controlled setting. Data was obtained from The Altered States Database, which contains psychometric data extracted from peer-reviewed articles published in MEDLINE-listed journals that used standardized and validated questionnaires. Our meta-analysis included data of the Altered States of Consciousness Rating Scale, the Mystical Experience Questionnaire (MEQ30), and the Hallucinogen Rating Scale (HRS). We used the Robust Variance Estimation Framework to obtain linear dose-response relationship estimates for each dimension of the given questionnaires. Ratings on most dimensions and subscales of the included questionnaires correlated positively with dose. Since subjective experiences are not only determined by dose, but also by individual differences and environmental factors, our results do not necessarily generalize to recreational use, as our analyses are based on data from controlled laboratory experiments. The paper at hand could serve as a general literature citation for the use of psilocybin in experimental and clinical research, especially for the comparison of expected and observed subjective drug experiences.

## INTRODUCTION

Psilocybin (4-phosphoryloxy-N,N-dimethyltryptamine) is a naturally-occurring tryptamine and the primary psychoactive component in *Psilocybe* mushrooms [1,2]. It has been used for centuries by native cultures for its ability to produce profound alterations of consciousness [3]. After ingestion, psilocybin is rapidly dephosphorylated to psilocin (4-hydroxy-N,N-dimethyltryptamine), which is mainly responsible for the psychedelic effects by agonist action at serotonin (5-HT) receptors, primarily the 5-HT2A receptor [4], similar to other classic psychedelics like Mescaline, Lysergic acid diethylamide (LSD), 5-Methoxy-N,N-dimethyltryptamine (5-MeO-DMT) and N,N-dimethyltryptamine (N,N-DMT) [5].

Even though the recreational use of psilocybin is widespread and psilocybin has found its applications in basic research as well as in clinical studies exploring therapeutic potentials [6–8], so far no dose-response relationships have been reported with regards to the psychoactive properties. Multiple reasons play a role in this. First, the doses applied in recreational use are often hard to quantify as the concentration of psilocybin in mushrooms is highly variable and typically not known to the consumer. Conducting laboratory studies with psilocybin is work and cost intensive as pharmacological intervention studies have to conform to strong security standards to ensure safety of participants [9]. Consequently, the amount of studies with controlled doses of psilocybin is limited. Finally, to establish dose-response relationships it is necessary that the response measure is accurately acquired across participants, which is challenging for psychological effects that depend on introspection [10].

The gold-standard in quantitative experimental research for measuring altered states of consciousness (ASC) experiences is the retrospect assessment with standardized and validated questionnaires [10–12]. Multiple questionnaires have been developed to quantify different aspects of ASC phenomena. Such psychometric measures allow direct comparisons between induction methods, individual’s responses, averaged group responses and different experimental settings.

Here we estimate dose-response relationships for orally administered psilocybin in healthy study participants based on the data from the Altered States Database [13], a collection of the currently available psychometric data on ASC experiences. Results of this analysis can be used to determine doses for experimental and clinical studies to induce the desired dose-specific subjective effects.

## METHODS

### Included data

We used the Altered States Database to identify peer-reviewed articles that contain suitable data for meta-analysis. The data was obtained from the Open Science framework repository in the version ASDB_v1.0_2019 (published in 2019-06) [14]. For the meta-analysis we only included datasets where the effects of orally administered psilocybin were investigated in healthy participants and any of the below specified questionnaires were applied. We therefore excluded the following datasets: Grob et al., Bogenschütz et al., Griffiths et al. [15–17] because these studies investigated the effects of psilocybin in patient populations; Griffiths et al. [18] because psilocybin administration was combined with meditation; Carhart-Harris et al. [19] because psilocybin was administered intravenously (i.v.) and it is unclear how the dose compares to oral administration. Three publications were duplicates, meaning data from the same experiment was reported (see **Table 1**). While one duplicate reported identical data [20], two duplicates differed in sample size [21,22]. Based on requests to the authors, we included the bigger samples [23,24].

**Table 1:** Summary of studies included in the meta-regression analysis. All studies were performed with healthy participants.

| Study | Sample Size | Study description | Data report | Psilocybin administration |
| --- | --- | --- | --- | --- |
| Umbricht 2002 [38] | N=18 | EEG with an auditory mismatch-negativity paradigm | 5D-ASC (converted) | Oral administration<br>Dosage:<br>(1) 280 µg/kg body weight |
| Hasler 2004 [94] | N=8 | Double-blind placebo-controlled within-subject design<br>EKG, blood pressure, body temperature and further questionnaire assessment | 5D-ASC | Oral administration as gelatin capsules<br>Dosages:<br>(1) 45 µg/kg body weight;<br>(2) 115 µg/kg body weight;<br>(3) 215 µg/kg body weight;<br>(4) 315 µg/kg body weight |
| Carter 2005a [95]<br>Wittmann 2007 [20] | N=12 | Double-blind placebo-controlled within-subject design<br>Binocular rivalry paradigm; time reproduction; auditory sensorimotor synchronization task; finger tapping; spatial span test | 5D-ASC | Oral administration as gelatin capsules<br>Dosages:<br>(1) 115 µg/kg body weight;<br>(2) 250 µg/kg body weight |
| Carter 2007 [23]<br>Carter 2005b [22] | N=10<br>(N=8) | Within-subject design with additional conditions: (2) Placebo; (3) 50mg Ketanserin; (4) Ketanserin + Psilocybin;<br>Multiple object tracking task; spatial working memory task | 5D-ASC | Oral administration as gelatin capsules<br>Dosage:<br>(1) 215 µg/kg body weight |
| Vollenweider 2007 [39] | N=16 | Double-blind placebo-controlled within-subject design<br>Prepulse-Inhibition of acoustic startle response | 5D-ASC (converted) | Oral administration as gelatin capsules<br>Dosages:<br>(1) 115 µg/kg body weight;<br>(2) 215 µg/kg body weight;<br>(3) 315 µg/kg body weight |
| Quednow 2012 [40] | N=16 | Double-blind within-subject design with additional conditions: (2) Placebo; (3) 40mg Ketanserin; (4) Ketanserin + Psilocybin;<br>Prepulse inhibition of acoustic startle response; color-word-stroop test | 5D-ASC (converted) | Oral administration as gelatin capsules<br>Dosage:<br>(1) 260 µg/kg body weight |
| Pokorny 2016 [96] | N=19<br>N=17 | Double-blind within-subject design with two groups (G1: N=19; G2: N=17) and additional conditions: (2) G1/G2: Placebo; (3) G1: 20mg Buspirone; or G2: 3mg Ergotamine; (4) G1: Buspirone + Psilocybin, or G2: Ergotamine + Psilocybin | 5D-ASC | Oral administration as gelatin capsules<br>Dosage:<br>(1) 170 µg/kg body weight |
| Schmidt 2012 [97] | N=20 | Double-blind within-subject design with additional conditions: (2) Placebo; (3) S-Ketamine<br>EEG with an auditory mismatch-negativity paradigm | 11-ASC | Oral administration<br>Dosage:<br>(1) 115 µg/kg body weight |
| Kometer 2012 [98] | N=17 | Double-blind placebo-controlled within-subject design<br>EEG with Facial Emotional Recognition task, Emotional Go/NoGo Task | 11-ASC | Oral administration as gelatin capsules<br>Dosage:<br>(1) 215 µg/kg body weight |
| Schmidt 2013 [99] | N=21 | Placebo-controlled<br>EEG; backward masking paradigm with facial affect discrimination | 11-ASC | Oral administration as gelatin capsules<br>Dosage:<br>(1) 115 µg/kg body weight |
| Bernasconi 2014 [100] | N=30 | Placebo-controlled<br>EEG; passive viewing of emotional face task | 11-ASC | Oral administration as gelatin capsules<br>Dosage:<br>(1) 170 µg/kg body weight |
| Carbonaro 2018 [76] | N=20 | Double-blind within-subject design with additional conditions: (4) 400mg/70kg DXM (5) placebo<br>Blood pressure, heart rate, pupil diameter, circular lights, balance, repeated administration of other questionnaires | 11-ASC<br>HRS<br>MEQ30 | Oral administration as gelatin capsules<br>Dosages:<br>(1) 143 µg/kg body weight (10mg/70kg);<br>(2) 286 µg/kg body weight (20mg/70kg);<br>(3) 429 µg/kg body weight (30mg/70kg) |
| Pokorny 2017 [24]<br>Preller 2016 [21] | N=33<br>(N=21) | Double-blind placebo-controlled within-subject design<br>Multifaceted empathy test, moral dilemma task (fMRI, Cyberball task) | 11-ASC | Oral administration as gelatin capsules<br>Dosage:<br>(1) 215 µg/kg body weight |
| Griffiths 2006 [101] (reported in Barret 2015 [33]) | N=30 | Double-blind, within-subject design with additional conditions (data from unblended control condition not included): (2) 40mg/70kg body weight<br>Methylphenidat<br>Group setting, meetings with monitor before and after sessions | HRS | Oral administration<br>Dosage:<br>(1) 429 µg/kg body weight (30mg/70kg) |
| Griffiths 2011 [102] | N=18 | Double-blind between-group crossover design, Descending or ascending dosage order | HRS<br>MEQ30 | Oral administration as gelatin capsules<br>Dosages: |
| (reported in Barret 2015 [33]) |  | Group setting, meetings with monitor before and after sessions |  | Placebo<br>(1) 71 µg/kg body weight (5mg/70kg);<br>(2) 143 µg/kg body weight (10mg/70kg)<br>(3) 286 µg/kg body weight (20mg/70kg);<br>(4) 429 µg/kg body weight (30mg/70kg) |
| Bravermanova 2018 [103] | N=20 | Double-blind, placebo-controlled EEG; Auditory mismatch negativity task | HRS | Oral administration as gelatin capsules<br>Dosage:<br>(1) 260 µg/kg body weight |
| Nicholas 2018 [104] | N=12 | Lying on sofa with eye-shades headphones and music;<br>Preparation of participants with guides;<br>Blood sample, ECG | MEQ30 | Oral administration as capsules<br>Dosages (given in escalating order):<br>(1) 300 µg/kg body weight;<br>(2) 450 µg/kg body weight;<br>(3) 600 µg/kg body weight |

If within-subject designs (repeated measures) were used, we supplemented the data dependency information to the datasets, based on the information in the original publications.

In an additional analysis, we included data on patients with alcohol dependence [16] and with cancer related psychiatric distress [15,17] on healthy participants to investigate the influence of pathologies on subjective experiences. The data in Griffith et al. [17] was given for a dosage range (22 or 30 mg per 70 kg) instead of a specific dose. However, since only 1 of 51 participants received 30 mg per 70 kg instead of 22 mg per 70 kg, we performed the additional analysis with the latter dose.

### Questionnaires

Our meta-analysis included data from three different questionnaires commonly applied in research on the subjective experiences induced by psychedelic substances: the Altered States of Consciousness Rating Scale, the Mystical Experience Questionnaire, and the Hallucinogen Rating Scale.

The Altered States of Consciousness Rating Scale [25,26] has become one of the most frequently used psychometric tools in the assessment of ASCs. It is supposed to investigate characteristics of ASCs that are invariant across various methods that are used to induced ASCs, including both pharmacological (e.g., psilocybin, ketamine, DMT, MDMA) and non-pharmacological methods (e.g., sensory deprivation, hypnosis, autogenic training). Over the course of more than 30 years, the questionnaire underwent several refinements finally leading to the currently used version which comprises 94 items [25,26]. Two different ways to analyze the questionnaire are in use. The first is referred to as *5D-ASC*, where the ratings of 66 items are combined to three core dimensions (1) *Oceanic Boundlessness*, (2) *Dread of Ego Dissolution* and (3) *Visionary Restructuralization*. Based on the remaining 28 items, the analysis is supplemented with two empirically derived scales which are considered specific to certain induction methods: (4) *Auditory Alterations* and (5) *Vigilance Reduction* [25,26]. The second way of analysis uses only 42 items of the three core dimensions [27] and summarizes the item scores along eleven factors. Correspondingly, these factors can be considered as subscales of the three core dimensions and have been termed: (1) *Experience of Unity*, (2) *Spiritual Experience*, (3) *Blissful State*, (4) *Insightfulness*, (5) *Disembodiment*, (6) *Impaired Control and Cognition*, (7) *Anxiety*, (8) *Complex Imagery*, (9) *Elementary Imagery*, (10) *Audio-Visual Synesthesia*, and (11) *Changed Meaning of Percepts*. Both analyses schemes have been validated and demonstrate good reliability (5D-ASC: Hoyt 0.88-0.95 [25,26]; 11-factorial structure: mean Cronbach’s alpha of 0.83 [27]).

The initial Mystical Experience Questionnaire (MEQ) was first used in the famous “Good Friday Experiment” [28,29], where it was intended to assess differences regarding aspects of mystical experience between a group taking the hallucinogen psilocybin and a control group taking a placebo. The items of the MEQ were chosen based on literature about mysticism including first-person accounts as well as theoretical work, most notably by James [30] and Stace [31]. The initial MEQ has been further developed, the most recent is a condensed version MEQ30 by MacLean et al. [32], consisting of 30 items and four empirical scales: (1) *Sacredness*, (2) *Positive Mood*, (3) *Transcendence of Time/Space*, and (4) *Ineffability*. This factor structure is currently recommended for analyses and has been assessed for reliability, yielding very good scores for all four subscales (Cronbach’s alpha: 0.80 to 0.95) [33,34].

Originally developed to quantify acute effects of synthetic dimethyltryptamine (DMT) [35], the Hallucinogen Rating Scale (HRS) has become a frequently used instrument in the assessment of hallucinogen induced ASCs. The initial construction of this questionnaire was based on systematic interviews with experienced hallucinogen users, describing the effects of smoked DMT freebase. The effects specifically induced by DMT, as well as general characteristic effects of hallucinogenic substances were intented to be covered by the resulting collection of items. The HRS measures six conceptually distinct dimensions of ASCs which were *a priori* defined and referred to as “clinical clusters”: (1) *Somaesthesia: interoceptive, visceral, and cutaneous/tactile effects, (2) Affect: emotional/affective responses*, (3) *Perception: visual, auditory, gustatory, and olfactory experiences, (4) Cognition: alterations in thought processes or content*, (5) *Volition: a change in capacity to willfully interact with oneself, the environment, or certain aspects of the experience*, and (6) *Intensity: the overall strength and course of the experience* [36]. Revision and refinement of the early versions finally resulted in the HRS 3.06 as the most recent version (available from the questionnaire’s author upon request), containing 100 statements, most of which are rated on a 5-point Likert scale. Reliability assessment indicates good internal consistency for the Affect, Somaesthesia, Cognition, and Perception scale [37].

### Standardization of data

Some studies reported the dose normalized to body weight (e.g. 25 mg per 70 kg body weight). To allow regressions-analyses, we converted the doses for all studies to μg (10^−6^ g) per kg body weight.

With regards to the psychometric data, some studies [20,38–40] reported the scores of the 5D-ASC as non-normalized sums of item scores. These scores were converted to the “percentage of maximum score”, in line with the Altered States Database [14].

### Statistical analyses

We performed linear meta-regression analyses for each dimension and subscale of the respective questionnaire. Although dose-response relationships are usually best described by a sigmoid function, the available data does not cover the upper and lower bounds to resemble a sigmoid function. We therefore modelled a linear dose-response relationship, which comprises a suitable approximation for the dynamic range of a sigmoid function. We used a random effects model to estimate the true underlying effects across studies [41], in terms of the intercept and slope of the dose-response function. The use of a random effects model accounts for between-study variance, which can result from e.g. differences in participant characteristics, interventions, the state of mind of participants (set), and the environment of drug intake (setting) across studies [42]. Multiple studies used a within-subject design, in which different doses were administered to the same set of study participants. To account for statistically dependent effect sizes, we used the Robust Variance Estimation Framework (RVE) developed by Hedges et al. [43] with small sample adjustment by Tipton [44]. The RVE method permits the inclusion of multiple effect size estimates from one study without the knowledge of the underlying covariance structure by assuming a common correlation p (0-1) between within-study effect sizes (p = 0.8 was used as the recommended default value [45]). To test if the choice of p had an effect on the obtained parameter estimates, we performed sensitivity analysis. The weights were calculated with the correlated effects model using the inverse of the sampling variance in combination with a method of moments estimator by Hedges et al. [43]. Heterogeneity was assessed by estimating the degree of inconsistency across studies with I^2^ [46,47] and the between-study variance with Tau^2^ [48]. Analyses were performed using the robumeta package [49] in R version 3.6.2 [50].

To allow comparability with previous reports, we used spiderplots for the visualization of the results for all questionnaires, inspired by reports of Vollenweider & Kometer [51], and Bayne et al. [52]. Spiderplots provide an overview of all questionnaire dimensions/subscale by showing the percentage of maximum score for different doses calculated with the linear regression estimates. Additionally, we present dose-response relationships for each dimension/subscale, including the effect sizes of the individual studies presented as circles. The size of a circle represents the magnitude of the calculated weight of a study sample. We generated spiderplots with the fmsb package [53] and scatterplots with the plot function in R version 3.6.2 [50].

## RESULTS

### Data description

We identified 20 publications with psilocybin administration in healthy study participants assessing psychometric data with validated questionnaires. Three reports were duplicate-reports on the same sample (See **Table 1**), which resulted in 7 samples of participants with 14 measurements (several studies include repeated measurements on the same sample) for the 5D-ASC (except for *Auditory Alterations* and *Vigilance Reduction* scales applied in 5 samples with 12 measurements), 5 samples with 7 measurements for the 11 subscale analysis of the ASC rating scale, 4 samples with 11 measurements for the MEQ30, and 3 samples with 8 measurements for the HRS.

### Dose-response relationships

Regression coefficients for the dose-response analyses and heterogeneity parameters are summarized in **Table 2**. Ratings on all dimensions and subscales of the included questionnaires correlated positively with psilocybin dose, except for the ASC rating scale subscales *Changed Meaning of Percept* and *Impaired Control and Cognition*. Spiderplots for each questionnaire and dose-response relationships for each dimension and subscale of the respective questionnaire are presented in **Figure 1** for 5D-ASC and the subscales of the ASC rating scale and **Figure 2** for MEQ30 and HRS.

**Table 2:** Meta-regression estimates for all included questionnaires with respective factors/dimensions/subscales. Coefficients (Coeff.) are presented with 95 % confidence intervals (CI) and standard errors (SE). The t-test statistic determines if a linear relationship exists under the null hypothesis that the slope is equal to zero. R2 represents the proportion of variability in subjective effects, which can be explained by psilocybin dose. Tau^2^ indicates the between-study variance and I^2^ indicates the degree of inconsistency across studies in percent. Intercepts’ estimates are rounded to the first decimal, except for the HRS due to its different range (0-4). Slope estimates are rounded to the third decimal considering its greater sensitivity to increasing dose.

| Outcome | Intercept |  |  | Slope |  |  | t (df) | p | R<sup>2</sup> | Tau<sup>2</sup> | I<sup>2</sup> |
| --- | --- | --- | --- | --- | --- | --- | --- | --- | --- | --- | --- |
|  | Coeff. | (95 % CI) | SE | Coeff. | (95 % CI) | SE |  |  |  |  |  |
| 5D-ASC |  |  |  |  |  |  |  |  |  |  |  |
| Auditory Alterations | 0.6 | (-8.7 – 9.9) | 1.71 | 0.043 | (0.006 – 0.081) | 0.0062 | 7.08 (1.5) | .040 | 0.97 | 4.21 | 22.30 |
| Oceanic Boundlessness | 4.0 | (-28.9 – 37.0) | 9.67 | 0.127 | (0.005 – 0.249) | 0.0388 | 3.28 (3.1) | .045 | 0.78 | 30.78 | 43.93 |
| Dread of Ego Dissolution | -2.2 | (-10.4 – 6.0) | 2.40 | 0.092 | (0.062 – 0.122) | 0.0093 | 9.81 (3.0) | .002 | 0.97 | 0.00 | 0.00 |
| Vigilance Reduction | 10.6 | (-6.4 – 27.6) | 4.15 | 0.098 | (-0.015 – 0.211) | 0.0252 | 3.88 (1.9) | .065 | 0.89 | 78.70 | 63.99 |
| Visionary Restructuralization | 6.3 | (-22.3 – 34.9) | 9.06 | 0.151 | (0.032 – 0.269) | 0.0390 | 3.86 (3.3) | .026 | 0.82 | 71.39 | 65.26 |
| 11 subscales of ASC rating scale |  |  |  |  |  |  |  |  |  |  |  |
| Anxiety | -0.7 | (-18.7 – 17.4) | 2.01 | 0.034 | (-0.059 – 0.125) | 0.0112 | 2.98 (1.2) | .166 | 0.88 | 0.00 | 0.00 |
| Audio Visual Synesthesia | 20.4 | (-55.7 – 96.4) | 11.70 | 0.053 | (-0.420 – 0.526) | 0.0580 | 0.92 (1.2) | .503 | 0.41 | 261.86 | 86.63 |
| Blissful State | 13.2 | (-34.4 – 60.7) | 7.50 | 0.115 | (-0.124 – 0.354) | 0.0321 | 3.57 (1.3) | .126 | 0.91 | 28.67 | 57.47 |
| Complex Imagery | 20.5 | (-99.8 – 140.8) | 16.94 | 0.116 | (-0.586 – 0.818) | 0.0860 | 1.35 (1.2) | .372 | 0.60 | 75.42 | 68.42 |
| Changed Meaning of Percepts | 32.7 | (4.1 – 61.3) | 4.40 | -0.015 | (-0.330 – 0.301) | 0.0370 | -0.39 (1.2) | .753 | -0.14 | 119.34 | 82.60 |
| Disembodiment | 11.4 | (-79.0 – 101.7) | 12.01 | 0.064 | (-0.452 – 0.581) | 0.0623 | 1.03 (1.2) | .463 | 0.46 | 39.53 | 55.89 |
| Elementary Imagery | 28.0 | (-133.0 – 189.0) | 23.26 | 0.103 | (-0.898 – 1.100) | 0.1200 | 0.86 (1.2) | .526 | 0.38 | 150.81 | 80.68 |
| Experience of Unity | 8.1 | (-0.6 – 16.8) | 1.03 | 0.097 | (0.041 – 0.153) | 0.0076 | 12.80 (1.3) | .025 | 0.99 | 0.00 | 0.00 |
| Insightfulness | 7.7 | (-33.4 – 48.9) | 6.49 | 0.104 | (-0.114 – 0.323) | 0.0290 | 3.60 (1.3) | .126 | 0.91 | 0.00 | 0.00 |
| Impaired Control & Cognition | 18.8 | (0.9 – 36.7) | 2.50 | -0.006 | (-0.119 – 0.108) | 0.0128 | -0.44 (1.2) | .725 | -0.20 | 25.21 | 67.03 |
| Spiritual Experience | -11.2 | (-59.5 – 37.2) | 6.92 | 0.150 | (-0.218 – 0.518) | 0.0475 | 3.16 (1.3) | .150 | 0.89 | 49.20 | 77.11 |
| MEQ30 |  |  |  |  |  |  |  |  |  |  |  |
| Ineffability | 48.2 | (-13.1 – 109.5) | 10.75 | 0.073 | (-0.020 – 0.166) | 0.0243 | 3.00 (2.3) | .081 | 0.80 | 45.28 | 59.02 |
| Mystical | 32.9 | (-28.5 – 94.3) | 9.90 | 0.081 | (-0.021 – 0.183) | 0.0264 | 3.07 (2.3) | .078 | 0.81 | 58.11 | 58.37 |
| Positive Mood | 49.9 | (2.7 – 97.0) | 7.97 | 0.058 | (-0.034 – 0.150) | 0.0239 | 2.42 (2.3) | .122 | 0.72 | 62.18 | 70.07 |
| Transcendence of Time & Space | 32.2 | (-22.5 – 86.8) | 9.23 | 0.090 | (0.000 – 0.180) | 0.0227 | 3.90 (2.3) | .048 | 0.87 | 77.49 | 71.95 |
| HRS |  |  |  |  |  |  |  |  |  |  |  |
| Affect | 1.24 | (-1.08 – 3.57) | 0.236 | 0.002 | (-0.001 – 0.005) | 0.0005 | 3.90 (1.7) | .078 | 0.90 | 0.01 | 41.39 |
| Cognition | 0.96 | (-1.56 – 3.48) | 0.261 | 0.003 | (-0.000 – 0.007) | 0.0007 | 4.53 (1.7) | .059 | 0.92 | 0.03 | 50.14 |
| Intensity | 1.96 | (1.58 – 2.34) | 0.040 | 0.002 | (0.001 – 0.003) | 0.0003 | 7.83 (1.7) | .024 | 0.97 | 0.02 | 50.25 |
| Perception | 0.88 | (-0.20 – 1.95) | 0.112 | 0.003 | (0.001 – 0.004) | 0.0003 | 10.59 (1.7) | .015 | 0.99 | 0.00 | 0.00 |
| Somaesthesia | 1.07 | (-1.94 – 4.08) | 0.313 | 0.002 | (-0.001 – 0.005) | 0.0006 | 3.11 (1.7) | .107 | 0.85 | 0.09 | 83.91 |
| Volition | 1.44 | (-0.23 – 3.11) | 0.154 | 0.001 | (-0.001 – 0.003) | 0.0004 | 2.21 (1.5) | .200 | 0.77 | 0.00 | 0.00 |

**Figure 1:**
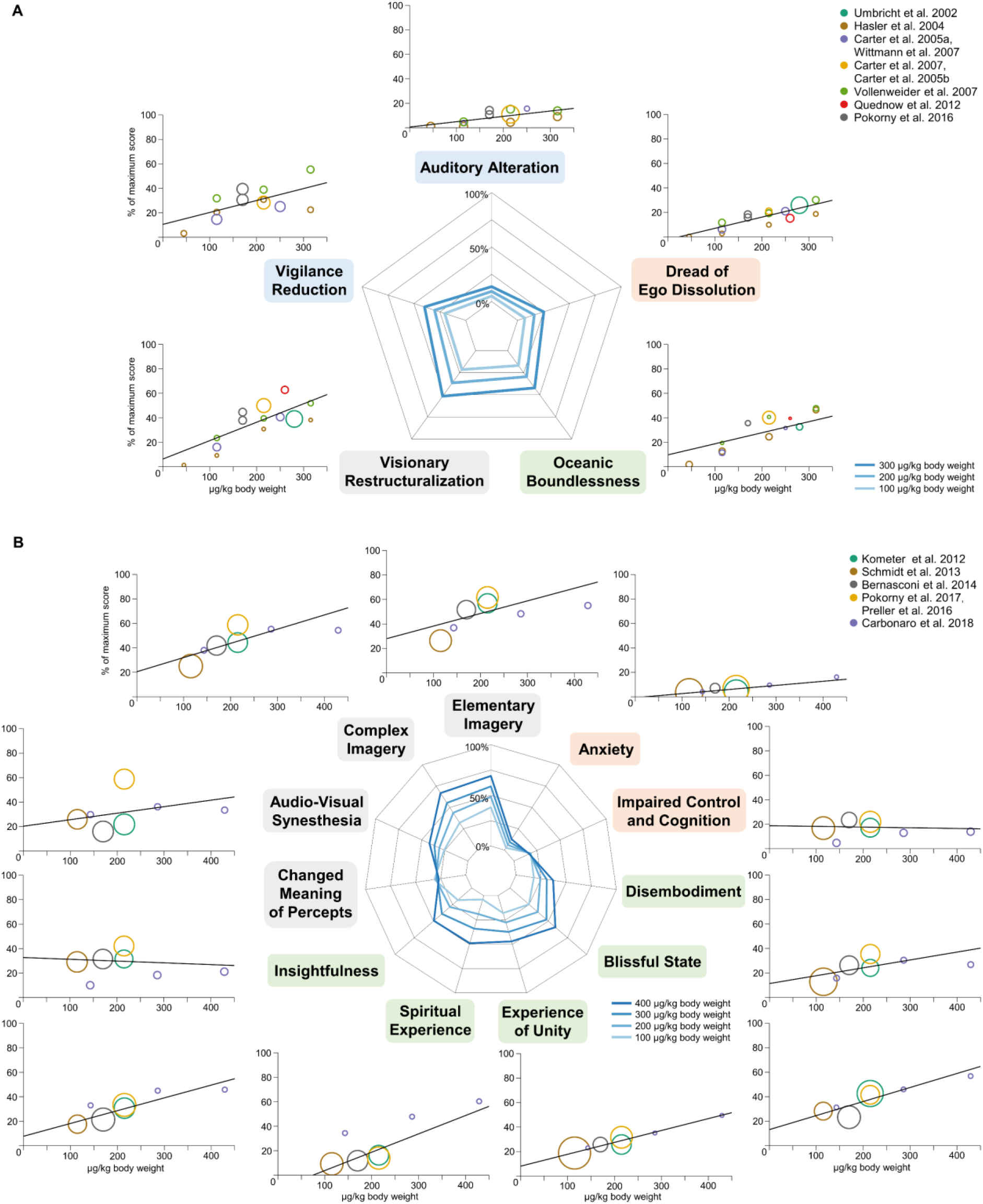
Dose-response relationships for the Altered States of Consciousness Rating Scale. Dose-specific subjective effects of psilocybin measured with the Altered States of Consciousness Rating scale. The data of this instrument can be analyzed according to a schema where items are organized into five factors, called “dimensions” of ASC experiences (5D-ASC) (See **A**). A finer grained quantification of specific aspects of subjective experiences is obtained when the questionnaire is analyzed according to an eleven factors schema (**B**). These eleven factors can be considered as subscales of the three core dimensions of the 5D-ASC, namely “Oceanic Boundlessness”, “Dread of Ego Dissolution” and “Visionary Restructuralization” (See corresponding coloring of the subscale names). Doses are given as μg per kg body weight; effects are given as percentage scored of the maximum score on each factor (questionnaire items are anchored by 0 % for “No, not more than usually” and 100 % for “Yes, much more than usually”). The color of the circles indicates data from the same sample of participants (same color corresponds to dependent data), while the circle size represents the weight of the data based on study variance (see Methods). Spiderplots were calculated with the regression coefficients for 100 - 300 μg/kg body weight on the 5D-ASC and 100 - 400 μg/kg body weight on the 11 subscales, corresponding to the range of doses which were included in the respective analysis. The color of individual scales corresponds to the primary dimensions and the respective subscales.

**Figure 2:**
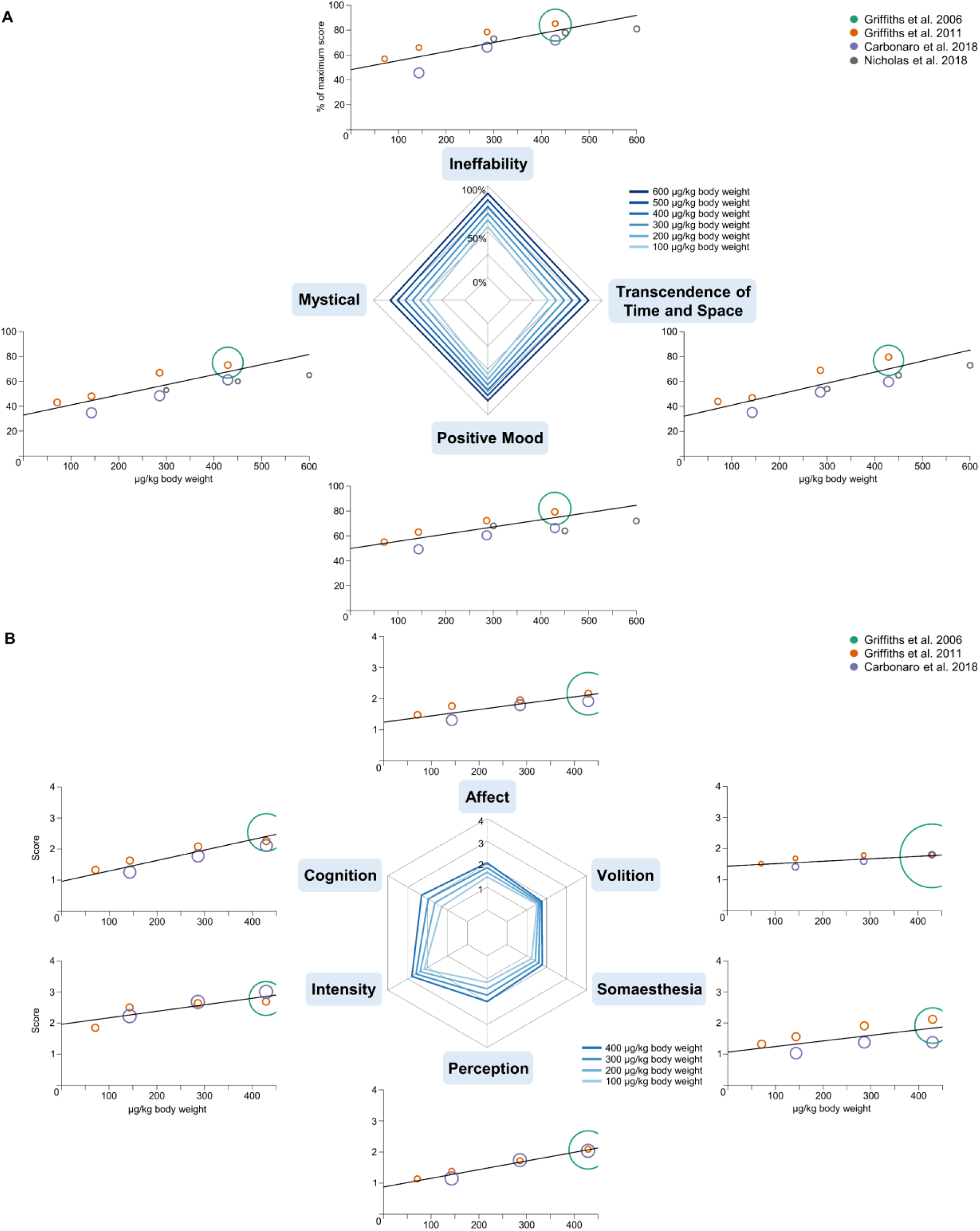
Dose-response relationships for MEQ30 and HRS. Dose-specific subjective effects of psilocybin for the psychometric instruments MEQ30 (**A**) and HRS (**B**). Doses are given as μg per kg body weight. Effects on the MEQ30 are presented as percentage scored on the maximum score. Effects on the HRS range from 0 – 4 (items in the questionnaire from 0 ”not at all” to 4 “extreme”). The color of the circles indicates data from the same sample of participants (same color corresponds to dependent data), the circle size represents the weight of the data based on study variance (see Methods). Spiderplots were calculated with the regression coefficients for 100 - 600 μg/kg body weight on the MEQ30 and 100 - 400 μg/kg body weight on the HRS, corresponding to the range of doses which were included in the respective analysis.

In the additional regression analysis, we included the available patient data on the 5D-ASC [15–17]. At the significance level of α = 0.05, we found no statistically significant interaction between patient status and dose (p > .05), indicating that subjective experiences of patients did not differ from healthy study participants. Correspondingly, the meta-regression estimates were comparable to the analyses on healthy study participants. Patient data increased the heterogeneity of the results (I^2^ and Tau^2^), except for the dimensions *Auditory Alterations* and *Vigilance Reduction* with comparable heterogeneity (see **Table S1**).

### Sensitivity analyses

To test the robustness of the estimated RVE parameters (intercept and slope), we examined if estimates were stable for different values of p (0-1) (See **Methods**). Across all analyses, intercept parameters differed only in the range of 0 - 0.18, and slope parameters were virtually identical, differing only in the range of 0 - 0.0007. Therefore, in line with Tipton [44], the sensitivity analyses produced robust effect size estimates for different values of p.

## DISCUSSION

Here we performed a meta-analysis on psychometric data to estimate linear dose-response relationships for psilocybin induced subjective experiences assessed with standardized questionnaires. Our analyses revealed positive correlations of effects and doses for almost all dimensions and subscales of the tested questionnaires.

For the 5D-ASC questionnaire, we found the strongest dose-responses for the scales *Visionary Restructuralization*, comprising alterations in perception, and *Oceanic Boundlessness*, comprising positively experienced ego dissolution, i.e. derealization and depersonalization associated with positive affect, ranging from heightened mood to euphoric exaltation. Interestingly, a medium dose-response was found for *Vigilance Reduction*, relating to states of drowsiness, reduced alertness and impaired cognitive function. Since classic psychedelics like psilocybin are usually characterized by a lack of sedation and clouding of consciousness, it was suggested that the effect of psilocybin on *Vigilance Reduction* rather reflect the state of contemplativeness, dreaminess and reduction of attentiveness [54]. *Dread of Ego Dissolution* associated with loss of self-control and anxiety exhibited a small dose-response with comparatively low rating scores. *Auditory Alterations*, relating to acoustic hallucinations and distortions in auditory experiences, were barely experienced. The analysis of the Altered States of Consciousness Rating scale along 11 subscales reflects finer facets of subjective experiences. Consistently with the 5D-ASC, the strongest dose-responses were found for subscales referring to *Visionary Restructuralization*, i.e. *Elementary* and *Complex Imagery*. In contrast, *Audio-Visual Synesthesia* and *Changed Meaning of Percepts* exhibited little modulation by dose. The intensity of the subscales referring to *Oceanic Boundlessness* increased with dose, especially for *Spiritual Experience* and *Blissful State*. In contrast, subscales referring to *Dread of Ego Dissolution* were barely modulated by dose and exhibited only small effects.

The MEQ30 questionnaire aims to measure different aspects of mystical-type experiences. The effects of psilocybin were characterized by relatively strong and equal effects on all four dimensions of the questionnaire. It had been suggested that scores of > 60 % on each of the four subscales indicate a complete mystical experience [33]. According to the obtained estimates, such experiences are expected for doses of approximately 350 μg/kg body weight and above. Interestingly, even very small doses caused relatively high scores, and the fitted regression lines have a relatively high y-axis intercept. While responses for the middle range of doses may be well predicted, the effects for doses in the lower range are not well covered by the given data.

On the HRS questionnaire, the dimensions *Cognition* and *Perception* showed the strongest dose response, whereas *Volition* was barely modulated by dose.

In summary, psilocybin mainly induced dose-dependent alterations in perception and positively experienced ego dissolution. Subjective experiences for high doses of psilocybin are characterized by all aspects of mystical-type experiences captured by the MEQ30 questionnaire. The given data does not support that higher doses of psilocybin would directly induce more aversive aspects of experiences, however, as the given data are average scores, it does not exclude that some individuals experiences highly challenging experiences.

### Variability of Subjective Experiences

Although psilocybin dose is the most important determinant of the acute psychedelic experience, there is considerable inter- and intra-individual variability in subjective responses to psilocybin [55–58]. To what extend responses to psilocybin varied among studies included in our main analysis is reflected in heterogeneity parameters. We assessed the between-study variance with Tau^2^ [48] and the degree of inconsistency across studies with F [46,47] (see **Table 2**). As expected, Tau^2^ estimates indicate variations between true effects for most of the analyzed dimensions. To what degree these variations are due to systematic differences between studies rather than random error is reflected by I^2^, given as percentage. Values for F are between 0 - 60 % for most analyses, indicating a small to moderate degree of inconsistency [59]. Effects were consistent (I^2^ = 0 %) for *Dread of Ego Dissolution* on the 5D-ASC, *Experience of Unity, Insightfulness* and *Anxiety* on the subscales of the ASC rating scale along with *Perception* and *Volition* on the HRS. In contrast, considerable inconsistencies (I^2^ > 75 %) were found for *Vigilance Reduction* on the 5D-ASC, and for the questionnaire subscales *Audio-Visual Synesthesia, Changed Meaning of Percepts, Elementary Imagery*, and *Spiritual Experience* as well as *Somaesthesia* on the HRS. For these scales, systematic differences between studies in terms of study population, study design or risk of bias accounted for more than 75 % of the observed variance. Despite the limited validity of the data along these scales, we produced robust dose-response relationships for the remaining scales, which allow for inferences on dose-specific subjective effects of psilocybin in a controlled setting.

Several non-pharmacological factors have been identified to cause variability in subjective experiences to psilocybin. A fundamental concept in psychedelic research is that the subjective experiences are highly dependent on the interaction of drug, *set* and *setting* [42,60,61]. *Set* constitutes the personality of the substance user and the preparation, expectation and intention of substance use [42,60]. In general, current mood and psychological distress prior to the psilocybin administration have been shown to contribute to the subjective experience [55,62,63]. Previous work demonstrated that the trait absorption and having clear intentions strongly correlated with all subjective experience measures [55,56]. Further, scoring high on trait extroversion was associated with increases in visionary experience [55,63,64]. Peak or mystical-type experiences were associated with the personality traits openness [63–65] and optimism toward life [64] as well as being in a state of surrender [57,58]. Likewise, feeling well prepared, having a recreational intention and emotional reappraisal reduced the occurrence of challenging experiences [56,64] which were associated with preoccupation [57,58] and emotional excitability prior to the experience [55]. While there are indications that challenging experiences are also associated with trait neuroticism [63,66,67], other studies found no such association [55,56,64]. These factors relating to *set* are often not obtained in studies but potentially contributed to the variability of subjective experiences. However, studies generally adhere to guidelines for the administration of psychedelics in a controlled experimental setting in order to minimize adverse effects [68,69], resulting in highly selected and well-prepared study populations.

The included studies also differed with regards to the applied *setting*, referring to the physical, social and cultural environment of drug administration [42,60]. While all included studies controlled the setting, the degree of interpersonal support, the amount and difficulty of tasks performed and the general ambient varied. Spatially confined neuroimaging settings were found to increase the likelihood of challenging experiences [55]. In our analysis, neuroimaging studies showed larger effect sizes on the respective dimensions and therefore might have led to overestimation of challenging experiences, although effect sizes were generally small.

Besides *set* and *setting*, other factors may also explain additional variability. Characteristics of the study populations mainly differed with regards to age and prior experience with psychedelics. Several studies reported that older study participants experience less *Impaired Control and Cognition* and tend to experience more of a *Blissful State* compared to younger study participants, whereas younger study participants more often report challenging experiences [55,62,63,70]. Hallucinogen-naïve study participants reported slightly more *Visionary Restructuralization, Disembodiment*, and *Changed Meaning of Percepts* compared to experienced psilocybin users [55,62]. In addition, inter-individual variability in pharmacokinetics was reported in terms of plasma psilocin levels and 5-HT2AR occupancy, which were found to correlate with the overall subjective experience [71–74]. Brain structure metrics could also exert influence, since the rostral anterior cingulate thickness correlated with subscales of the Altered States of Consciousness Rating Scale *Oceanic Boundlessness* [75]. Nevertheless, it should be noted that studies which used linear regression models to predict the acute subjective experience reported relatively large proportions of unexplained variances in subjective responses to psilocybin [55,56]. Despite psilocybin dose being the most important determinant of subjective experiences, the influence of non-pharmacological factors should be considered when interpreting dose-response relationships.

### Comparison to previous reports

Our results demonstrate that psilocybin intensified almost all characteristics of ASCs that are measured by the given questionnaires in an approximately linear dose-dependent manner. While Vollenweider & Kometer [51] and Studerus et al. [54] suggested an approximately linear relationship on data assessed within their research group, we extended this work by including all available psychometric data across studies. In line with our results, the strongest dose-responses were found for perceptual alterations, followed by subscales relating to *Oceanic Boundlessness*. While they report dose-dependent increases of effect sizes for *Changed Meaning of Percepts* and *Impaired Control and Cognition*, our analysis revealed weak effects with considerable inconsistencies, potentially due to methodological differences between studies with regards to the amount and difficulty of tasks performed during the psilocybin-induced state. In Studerus et al. [54] and in most studies in our analysis, participants engaged in performing tasks for a considerable amount of time, whereas Carbonaro et al. [76] is the only study in which participants received several doses and were encouraged to focus on the phenomenology. *Changed Meaning of Percepts* and *Impaired Control and Cognition* might have been more apparent with increased interaction with researchers compared to minimum interactions. This could also explain why *Audio-Visual Synesthesia* showed a stronger dose-response and *Spiritual Experience* was less dose-dependent compared to our results.

Other factors like induction method, health status and additional interventions could also affect subjective experiences. Since we only included studies where psilocybin was administered orally, we excluded one study with intravenous administration of 1.5 mg and 2 mg [19]. Despite the considerably smaller dose quantities, the pattern of responses on the 5D-ASC were roughly comparable to our results, corresponding to a low oral dose (i.e. 100-150 ug/kg body weight) for 1.5 mg intravenously and a medium oral dose (i.e. 200-250 ug/kg body weight) for 2 mg intravenously. In two studies, psilocybin was administered either in combination with spiritual practice support [18] or to experienced meditators [77]. Compared to our analysis, both studies reported a similar pattern of responses for visual and auditory alterations for the administered doses, but substantially larger effect sizes for *Oceanic Boundlessness* and smaller effect sizes for *Dread of Ego Dissolution* than predicted by our dose-response estimates. Moreover, when we included data from patients with alcohol dependence disorder [16] and life-threatening cancer [15,17] in the 5D-ASC analysis, dose-response estimates were relatively similar, but heterogeneity increased for most dimensions. Effect sizes for *Oceanic Boundlessness* were slightly higher, while effect sizes for *Dread of Ego Dissolution* were slightly lower, whereas visual and auditory alterations did not differ. This is not surprising, since these studies were designed to facilitate peak or mystical-type experiences, which were shown to correlate with positive long-term outcomes [16,17,65,78–85].

Other classic psychedelics may induce comparable subjective experiences to psilocybin, considering that participants in early studies failed to discriminate between the subjective experiences of psilocybin, LSD and mescaline [86,87]. So far, no dose-response relationships have been established for those substances, but a report by Liechti [88] analyzed data on LSD from three studies of their research group, which showed a comparable pattern of dose-responses on the Altered States of Consciousness Rating Scale, and MEQ30 questionnaires. Whereas the scales *Audio-visual Synesthesia* and *Changed Meaning of Percepts* exhibited a strong dose-response with LSD, these dimensions where barely dose-dependent in our analysis. The *Mystical* dimension on the MEQ30 was also less dose-dependent and showed smaller effect sizes compared to our results. The comparison is, however, limited by the small amount of psychometric data for LSD. Studies with N,N-DMT and 5-MeO-DMT report largely comparable pattern of response to psilocybin on the MEQ30 and HRS [36,89,90]. Despite the reported commonalities in subjective experiences to other classic psychedelics, the characterization of differences requires the utilization of standardized questionnaires in future research in order to establish dose-response profiles, which could then serve as a general reference to compare and infer subjective experiences.

### Limitations

Results of the present study need to be understood in relation to the method-immanent limitations. First, dose-response relationships are, like most biological processes, typically sigmoidal relationships. Since the available data does not cover the upper and lower bounds to resemble a sigmoid function, we modelled a linear function to approximate the dynamic range of a sigmoid function. Our results indicate approximately linear dose-response relationships, which is consistent with previous reports [51,54]. However, finding relatively high y-axis intercepts for multiple questionnaire scores indicate that at least the range of low and very low doses (e.g. microdosing) cannot be predicted well by the obtained models. Second, the statistical method RVE permits the inclusion of statistically dependent effect sizes to estimate reliable meta-regression estimates, but it is not intended to provide precise variance parameter estimates, nor test null hypotheses regarding heterogeneity parameters [91]. Third, generalizability of our results is limited due to the amount of available studies with generally small sample sizes in a controlled setting. Recruitment strategies were prone to self-selection bias and inclusion criteria resulted in highly selective study populations. The obtained results do not necessarily apply to the general population or to psilocybin administration in a recreational or less controlled setting. Lastly, the study of subjective experiences depends on introspection and is more challenging to assess than other physiological parameters, especially because these experiences often go beyond the previously experienced epistemic range [92,93].

In conclusion, the subjective experience induced by psilocybin is mainly characterized by perceptual alterations and positively experienced ego dissolution with the ability to occasion mystical-type experiences. Although the psychedelic experience is also dependent on non-pharmacological variables, we established robust dose-response relationships for most dimensions, which can be used as a general reference for relating expected and observed dose-specific effect. The results not necessarily generalize to recreational use, as our analyses were based on data from controlled laboratory experiments in healthy, highly selected study participants. Future research should facilitate comparison of subjective experiences by utilizing standardized questionnaires in order to improve dose-response profiles and inform future clinical studies.

## Acknowledgement

We thank the students of the interdisciplinary MSc course “How it feels to be on....?”, held at the Institute of Cognitive Science, Universität Osnabrück for contributions to the Altered States Database and establishing the applied meta-analytic approach, in particular Greta Häberle, Carolin Gaß, Geeske Sieckmann, Regina Gerber, Marcel Lommerzheim, Shadi Derakhshan, Felix Beering, Kimberly Gerbaulet, Lukas Niehaus, Andreas Strube and Aaron Gutknecht. We thank Gabriele Inciuraite for proofreading and helpful suggestions on the manuscript.

