## Supplementary material for "How does it feel to be on psilocybin? Dose-response relationships of subjective experiences in humans": Table S1

**Appendix**

Results for the additional meta-regression analysis, conducted like the main analysis but including data of patients. Included data consists of a study on patients with alcohol dependence [1] and two studies on patients with psychological distress due to life-threatening cancer [2,3].

|  |  |  | Intercept | | |  | Slope | | |  |  |  |  |  |  |  |
| --- | --- | --- | --- | --- | --- | --- | --- | --- | --- | --- | --- | --- | --- | --- | --- | --- |
|  | Outcome |  | Coeff. | (95 % CI) | SE |  | Coeff. | (95 % CI) | SE |  | t (df) | p | R² |  | Tau² | I² |
| 5D-ASC | |  |  |  |  |  |  |  |  |  |  |  |  |  |  |  |
|  | Auditory Alterations |  | 3.1 | (-9.3 − 15.5) | 7.11 |  | 0.035 | (-0.011 − 0.081) | 0.0086 |  | 4.06 (1.7) | .076 | 0.91 |  | 3.65 | 20.83 |
|  | Dread of Ego Dissolution |  | 0.5 | (-17.4 − 18.4) | 4.98 |  | 0.071 | (-0.010 − 0.153) | 0.0229 |  | 3.12 (2.5) | .067 | 0.80 |  | 53.12 | 19.25 |
|  | Oceanic Boundlessness |  | 10.0 | (-14.4 − 34.4) | 7.74 |  | 0.126 | (0.036 − 0.215) | 0.0293 |  | 4.29 (3.2) | .020 | 0.85 |  | 74.85 | 109.22 |
|  | Vigilance Reduction |  | 15.2 | (-9.7 − 40.2) | 6.78 |  | 0.067 | (-0.066 − 0.200) | 0.0342 |  | 1.96 (2.2) | .175 | 0.64 |  | 61.76 | 53.42 |
|  | Visionary Restructuralization |  | 8.1 | (-14.4 − 30.6) | 7.11 |  | 0.153 | (0.056 − 0.250) | 0.0312 |  | 4.90 (3.2) | .015 | 0.88 |  | 67.75 | 72.49 |

**Table S1**: Meta-regression estimates for the 5D-ASC questionnaire with respective dimensions. Intercepts estimates are rounded to the first decimal. Slope estimates are rounded to the third decimal considering its greater sensitivity to increasing dose. Coefficients (coeff.) are presented with 95 % confidence intervals (CI) and standard errors (SE). I² indicates the degree of inconsistency across studies in percent and Tau² the between-study variance.
